# Simulation with RADinitio Improves RADseq Experimental Design and Sheds Light on Sources of Missing Data

**DOI:** 10.1101/775239

**Authors:** Angel G. Rivera-Colón, Nicolas C. Rochette, Julian M. Catchen

## Abstract

Restriction-site Associated DNA sequencing (RADseq) has become a powerful and versatile tool in modern population genomics, enabling large-scale genomic analyses in otherwise inaccessible biological systems. With its widespread use, different variants on the protocol have been developed to suit specific experimental needs. Researchers face the challenge of choosing the optimal molecular and sequencing protocols for their experimental design, an often-complicated process. Strategic errors can lead to improper data generation that has reduced power to answer biological questions. Here we present *RADinitio*, simulation software for the selection and optimization of RADseq experiments via the generation of sequencing data that behaves similarly to empirical sources. *RADinitio* provides an evolutionary simulation of populations, implementation of various RADseq protocols with customizable parameters, and thorough assessment of missing data. Using the software, we test its efficacy using different RAD protocols across several organisms, highlighting the importance of protocol selection on the magnitude and quality of data acquired. Additionally, we test the effects of RAD library preparation and sequencing on allelic dropout, observing that library preparation and sequencing often contributes more to missing alleles than population-level variation.

## Introduction

Type II restriction enzymes have powered population genomic experiments since their identification in the 1970s (Kelly & Smith, 1970; Smith & Welcox, 1970) being coupled with each new generation of technology to provide more markers, at lower costs (Schlötterer, 2004). Restriction site-Associated DNA sequencing (RADseq), which combined restriction enzymes with massively parallel, short-read sequencing, has further democratized population genomics (Catchen et al., 2017). RADseq has enabled numerous ecological and conservation genomics studies and has allowed for the development of genomics resources in non-model systems which were previously thought to be inaccessible and cost prohibitive (Andrews, Good, Miller, Luikart, & Hohenlohe, 2016; Narum, Buerkle, Davey, Miller, & Hohenlohe, 2013). Since the inception of the original single-digest RADseq (sdRAD) protocol (Baird et al., 2008; Etter, Bassham, Hohenlohe, Johnson, & Cresko, 2011), other variants of reduced representation libraries employing restriction enzyme cutsites for the selection of genetic markers have been developed, including GBS (Elshire et al., 2011), double-digest RADseq (ddRAD) (Peterson, Weber, Kay, Fisher, & Hoekstra, 2012), RAD capture and Best RAD (Ali et al., 2016), 2b-RAD (Wang, Meyer, McKay, & Matz, 2012), ezRAD (Toonen et al., 2013), RADcap (Hoffberg et al., 2016), and FecalSeq (Chiou & Bergey, 2018). This family of protocols, referred hereafter as RADseq, while displaying individual benefits and disadvantages, have been designed for specific experimental contexts, and are a testament of the applicability of the general molecular approach. Independent of the specific molecular protocol used, RADseq has proven to be a versatile technique in a variety of genomics contexts, including the generation of linkage maps (Amores, Catchen, Ferrara, Fontenot, & Postlethwait, 2011; Amores, Wilson, Allard, Detrich, & Postlethwait, 2017; Small et al., 2016), *de novo* population genomics (Jeffery et al., 2017; Portnoy et al., 2015), landscape genomics (Bay et al., 2018; Dudaniec, Yong, Lancaster, Svensson, & Hansson, 2018), reference-based genome scans (Bassham, Catchen, Lescak, von Hippel, & Cresko, 2018), and phylogenomics/phylogeography (Cristofari et al., 2016; Razkin et al., 2016; Suchan et al., 2017).

Alongside its widespread use as a molecular protocol, a variety of bioinformatic software has been designed to work specifically with RADseq data (Catchen et al., 2011; Catchen, Hohenlohe, Bassham, Amores, & Cresko, 2013; Chong, Ruan, & Wu, 2012; Eaton, 2014; Melo & Hale, 2019; Puritz, Hollenbeck, & Gold, 2014; Rochette, Rivera-Colón, & Catchen, 2019), and methods have been developed to optimize the application of these software after data generation (Ilut, Nydam, & Hare, 2014; McCartney-Melstad, Gidiş, & Shaffer, 2019; Paris, Stevens, & Catchen, 2017; Rochette & Catchen, 2017). However, software and parameter optimization protocols are not effective if the underlying sequenced data has captured little of the true biological signal. Error in RADseq data can be traced to two non-mutually exclusive sources: 1) the unsuitable choice and application of a molecular protocol, and 2) problems with library preparation and sequencing. Regarding the selection of molecular protocol, it has been noted that different techniques are often chosen based on their popularity and experimental simplicity, potentially disregarding the suitability of different RADseq methods for different species and experimental designs (Andrews et al., 2016; Campbell, Brunet, Dupuis, & Sperling, 2018). In addition, we observe that a relatively high number of experiments are being conducted without a sufficient assessment of the biological system, resulting in less than optimal library preparation, high missing data, and low coverage, leading to poor analytical results. Independent of the cause, the generation of low-quality data wastes efforts and resources, and therefore, it is in a researcher’s best interests to select and apply a molecular protocol that yields the best data possible.

The likelihood of experimental success can be significantly increased with *prospective* data simulation, allowing for the exploration of simulated data generated under different experimental conditions, before sample collection, library preparation, and sequencing. These simulations would allow researchers to test the behavior of different molecular protocols in their systems, as well as assessing the magnitude of data recovered given variable experimental conditions. While tools exist for the simulation of RADseq data, limitations in those tools prevent a complete assessment of the properties of sequences generated. *Simrlls* (Eaton, Spriggs, Park, & Donoghue, 2016) is a pipeline optimized for the simulation of phylogenomic data at various evolutionary scales but generates random sequence data that is highly artificial, with no relation to the species and genome of interest, and without any effects of library preparation and sequencing. *simRAD* (Lepais & Weir, 2014) can generate data from existing reference sequencing, allowing to test for cutsite frequencies under different RADseq protocols, but it does so for a single sample without accounting for any genetic variation across individuals. *ddRADseqtools* (Mora-Márquez, García-Olivares, Emerson, & López de Heredia, 2017) can generate ddRAD data both from a reference sequence and via random sequence generation. It can simulate multiple library configurations, such as differences in insert size, adapter, PCR duplicates and coverage. While it also simulates variants for multiple samples, these are independent across individuals, and have only fixed effects over allele dropout. Finally, *ddrage* (Timm, Weigand, Weiss, Leese, & Rahmann, 2018) simulates ddRADseq data under multiple library configurations, in both a *de novo* and reference context. It can also generate variants that are shared across individuals. However, the modeling of allele dropout due to variants, and the effects of PCR duplicates, are generated from fixed probabilities and do not respond dynamically to the behavior of the variants and the library preparation process. Aside from library preparation parameters, none of the available software can generate variants that are relevant for population genetics, meaning that they limit the capacity of testing RADseq protocols in a biologically meaningful context.

Here, we present *RADinitio*, a software for the assessment of RADseq experiments via the generation of simulated data. Our software generates phased variants based on a coalescent simulation (Kelleher, Etheridge, & McVean, 2016) under a user-defined demographic model, which ensures that variants are generated and shared across individuals. Instead of simulating within a RAD locus context, variants are generated genome-wide and thus the library is generated within the constraints of an existing genotype pool, more accurately simulating downstream effects like allelic dropout. *RADinitio* also simulates multiple components of the RADseq library preparation process – including protocol type, restriction enzymes, fragment size selection, library amplification, and sequencing coverage. Metadata is generated across different parts of the pipeline, which ensures the traceability of the simulation process and enables downstream comparisons. In addition to the *prospective* simulation of data, *RADinitio* allows for *retrospective* simulations, where information from existing empirical datasets are used for *de novo* simulations for comparative purposes.

Using *RADinitio*, we explore previously discussed components of RAD library preparation, such as the selection of a single or double-digest protocol, enzyme selection in a known reference genome, and RAD loci estimation with an unavailable reference. Additionally, we explore sources of error in RADseq data, and how problems in library preparation and sequencing likely play a larger role in allele dropout than genetic polymorphism in a study population. *RADinitio* allows researchers to fully explore possible variables in their data generation process to ensure that their protocol selection and components of their library preparation process is performed optimally, within the limitations of technical and experimental error.

## Materials and Methods

### RADinitio pipeline

#### 1. Simulating a base metapopulation

Generation of genetic variants in *RADinitio* is based on the demographic models available in the *msprime* simulation software (Kelleher et al., 2016). *msprime* generates abstract genetic variants via the simulation of coalescent trees for various samples given a user-defined neutral demographic model, which can later be converted into individual haplotypes. For *RADinitio*, we generate one *msprime* coalescent simulation for each chromosome/scaffold available in a reference sequence provided by the user (in FASTA format), using the length of this sequence as the *msprime* input `length` value (Fig. 1A). By default, *RADinitio* provides *msprime* with a simple island model (Fig. 1B) defined as a number of populations, each with a defined effective population size *N*_*e*_. Each population has a symmetrical per-generation migration rate. Other simulation parameters, including the mutation and recombination rates can also defined by the user. In fact, the full complexity of *msprime* can be invoked while keeping compatibility with *RADinitio*. Even the simulation of non-neutral evolutionary events, such as selective sweeps (Haller, Galloway, Kelleher, Messer, & Ralph, 2019; Haller & Messer, 2019), can be compatible with *RADinitio* (Fig. S1).

**Figure 1.**
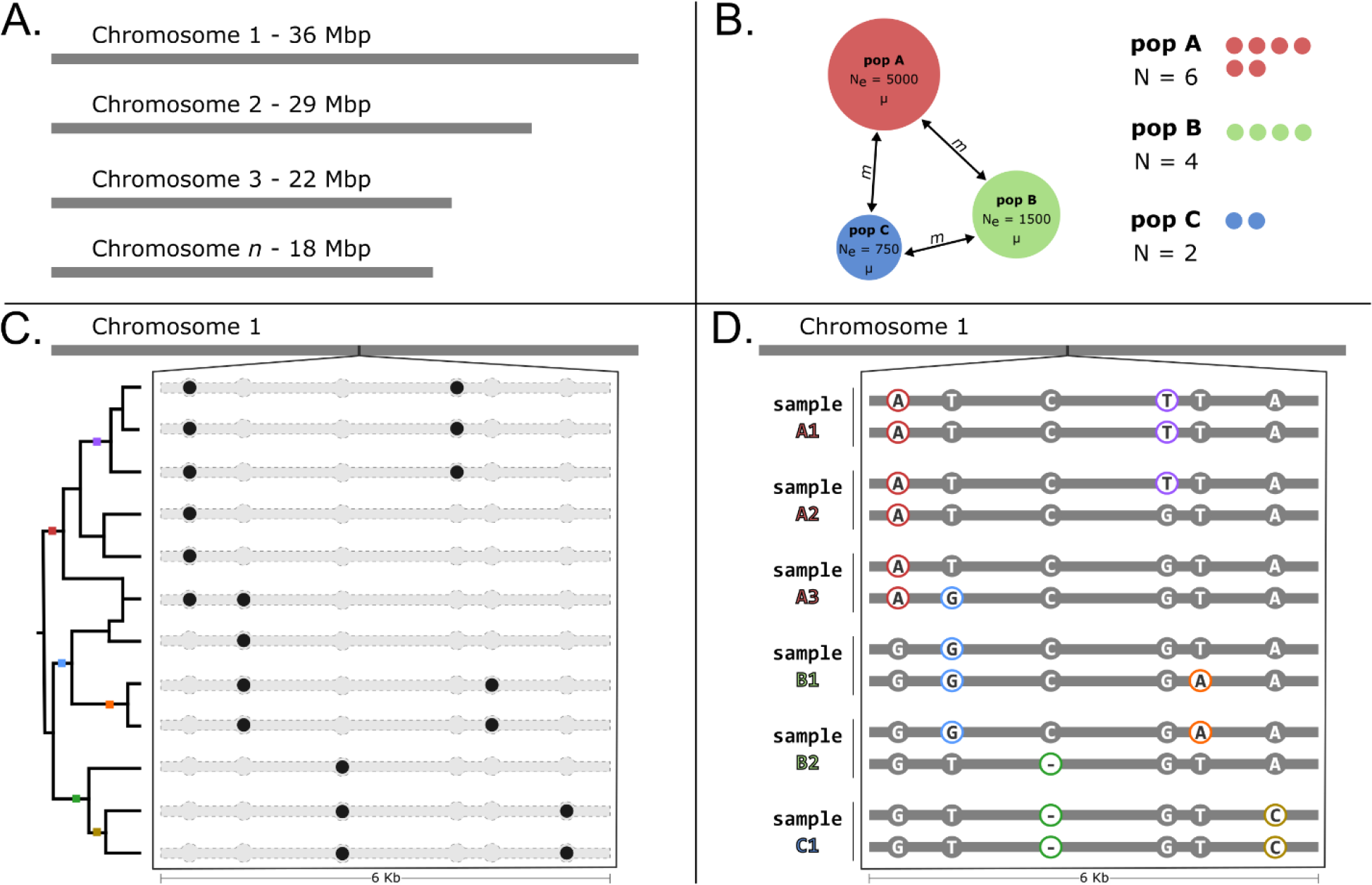
RADinitio simulates genetic variation over a reference sequence under the constrains of a demographic model. (A) A reference genome serves as basis for simulation space and future extraction of sequences. (B) Using *msprime*, we simulate three populations (A, B, and C) of different sizes (*N*_*e*_), with a symmetric migration rate *m*, from which we sample chromosomes which will later become diploid individuals. (C) We observe the tree topology, mutations, and genotype matrix for a 6Kb regions in a hypothetical chromosome. The variants (black circles) are the product of coalescent simulation with recombination of our defined metapopulation over which mutations are added at a rate *μ* (colored squares). (D) The mutations generated in (C) are converted to alleles by comparing the genotype matrix against the reference sequence. After obtaining reference alleles from the reference genome (gray circles), alternative alleles (colored circles) are generated accordingly.

The result of the *msprime* coalescent simulations is a per-chromosome/scaffold genotype matrix stored as a VCF file (Danecek et al., 2011) (Fig. 1C). These generated VCFs contain phased genotypes, but they are represented uniformly by the digits ‘0’ and ‘1’, not the actual identity (alleles) of the individual variants. *RADinitio* obtains the final genotypes by projecting the genotype matrix back to the reference sequence, associating reference alleles and generating alternative alleles (Fig. 1D). The alternative alleles are obtained by comparing the reference alleles against a user-defined substitution model, which can include indels. By default, this model has equal substitution probability for all non-reference nucleotides, with a fixed indel probability with length sampled from a Poisson distribution with *λ*=1. After processing, variants are then merged into a single genome-wide VCF file containing all simulated variants.

The use of this coalescent simulation method for the generation of variants allows for the systematic generation of individual haplotypes that follow the behavior of those observed in natural populations. Haplotypes across individuals will be related based on the underlying demographic constraints, and because of the presence of recombination, haplotypes behave as discrete elements in the genome. By simulating mutations in a genome-wide context instead of a RAD locus context, we can then observe the effects of shared variants in the downstream sampling of loci. This process mimics the real-life scenario in which a library is prepared and sequenced under the constraints of an existing pool of genotypes, with polymorphic variants impacting the dropout of alleles (via mutations in a restriction enzyme cutsite sequences) in individuals within a related population. This simulated population can then be *in silico* sequenced by *RADinitio* using libraries generated under different configurations of an evolutionary model.

#### 2. Defining RAD loci

Reference RAD loci are extracted via *in silico* digestion of the reference genome chosen by the user (Fig. 2A). *RADinitio* first iterates over all sequence in the reference genome, storing the position of each cutsite for the set of user-specified restriction enzymes. The raw cutsite positions are filtered to remove closely overlapping loci – based on a default locus length of 1 kilobase (Kb), to simulate the effects of shearing biases on small DNA fragments that are the product of close restriction cutsites (Davey et al., 2013), as well as loci close to the ends of scaffolds or chromosomes. A tally of the total number of loci and their position in the genome, including those marked as removed, is generated as an output table. For each kept cutsite, sequences 1Kb upstream and downstream are extracted and stored (in a FASTA file) as two separate tags, representing the reference state of all RAD loci and serving as a template for the downstream generation of alleles and sequencing reads.

**Figure 2.**
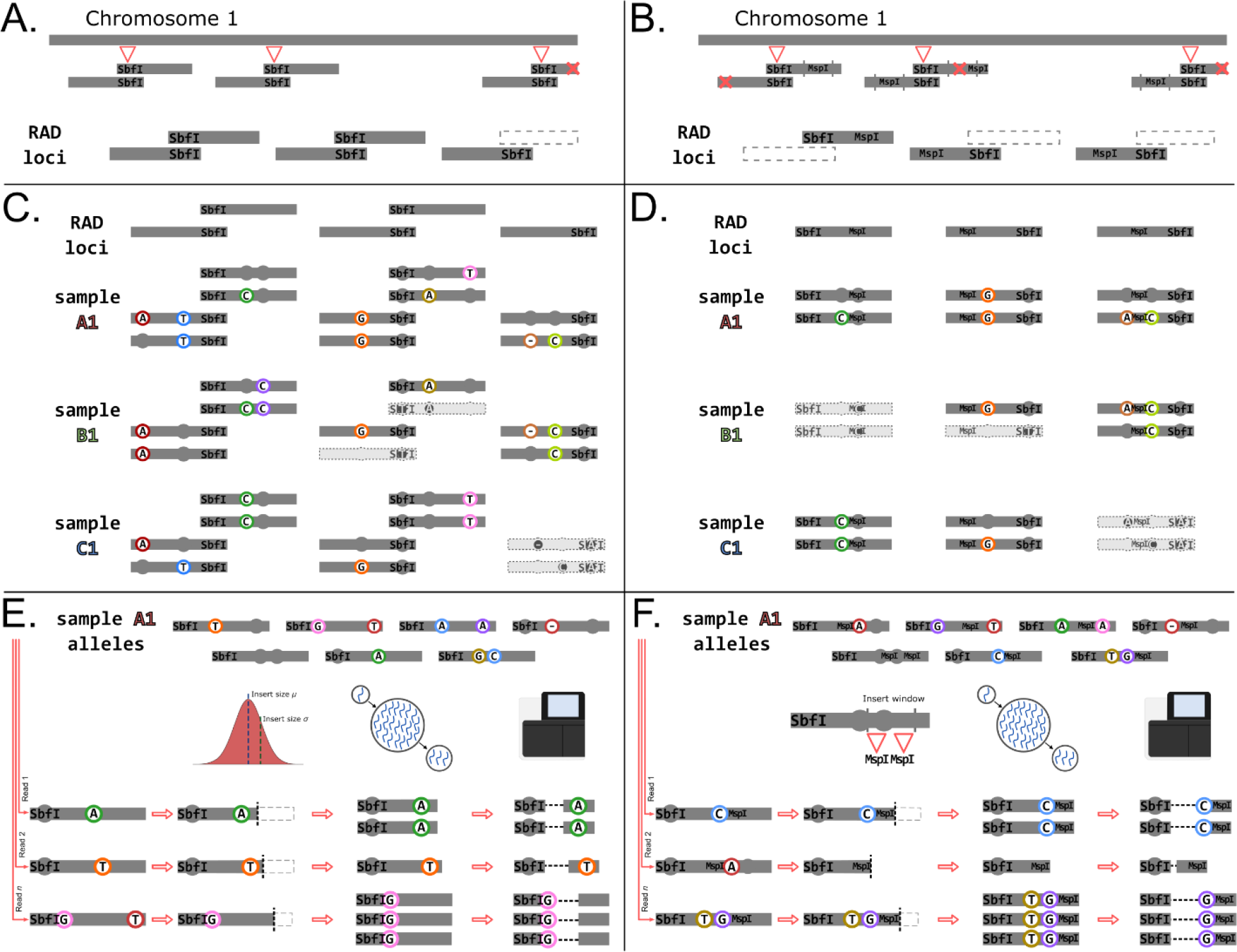
RADinitio simulates library preparation and sequencing. (A) Single-digest RAD loci are obtained by *in silico* digest of the reference sequence with a restriction enzyme, e.g. *SbfI*. Each locus is extracted and potentially filtered if overlapping with adjacent loci or sequence boundaries. (B) Double-digest RAD loci are extracted via *in silico* digest using a pair of enzymes, e.g. *SbfI* and *MspI*, and removed if lacking restriction sites within the specified insert size range. (C) Simulated mutations (Fig. 1) are incorporated into sdRAD loci to generate alleles for each sample. An allele is dropped if its restriction cutsite sequence has been mutated (rectangles with dashed outlines). (D) When generating alleles in ddRAD, sequences for both restriction cutsites are checked, dropping those in which it has been modified for either enzyme. (E) For sdRAD, alleles are sampled to simulate sequencing reads. Insert lengths for each molecule are sampled from a normal distribution, mimicking mechanical shearing. Following *in silico* amplification, which can potentially lead to PCR duplicates, each molecule is then independently sequenced. (F) The same *in silico* PCR and sequencing happens for ddRAD, but insert sizes are obtained from digestion with the frequent-cutter enzyme.

In the case of a ddRAD library (Fig. 2B), the user has the option to specify the type of size selection to employ within the *in silico* library, since the insert length of the library (which is defined during size selection) will affect which loci are recovered for amplification and sequencing (larger insert lengths result in more ddRAD loci). *RADinitio* first stores and filters the positions of all forward cutsites, similar to the simulation of single-digest libraries. The extracted reference sequences are then scanned for the reverse cutsite, coming from the second, frequent-cutting restriction enzyme. Only cutsites found within the range of the user defined insert size distribution, which is 150bp wide by default, are kept. Reference sequences without reverse cutsites, or with cutsites outside the insert size range, are discarded, and a tally of all loci, both kept and discarded, is generated. The sequence for surviving loci is truncated to the length of the distal-most reverse cutsite, allowing for the selection of different 3’ cutsites across different reads coming from the same reference locus

#### 3. Per-individual RAD alleles

The genome-wide set of polymorphisms (SNPs and indels) generated by *msprime* are intersected with the extracted RAD loci so as to keep only variants present within RAD loci, reducing the amount of downstream processing. For each sample, *RADinitio* iterates over all RAD loci, selecting the corresponding sample variants for each RAD allele (Fig. 2C,D). The reference locus sequence is then modified to include the corresponding allelic variants. Only sequences with intact cutsites (containing no mutations within the restriction enzyme recognition sequences) are saved, simulating the allele dropout process. The program outputs per-sample FASTA files containing the modified allele sequences for each RAD locus, and a report of the status of all RAD alleles across all samples.

#### 4. Library amplification and sequencing

Paired-end reads are generated from the extracted RAD alleles after simulated library preparation and sequencing (Fig. 2E,F). This *in silico* enrichment and sequencing begins by generating a PCR duplicate/error distribution (Rochette & Rivera-Colón, *in prep*). Briefly, this distribution is the result of modeling three steps that represent 1) the pre-amplification library complexity as the number of unique template molecules, 2) the complexity of the amplified library in frequencies of the amplified clones (sets of molecules originating from a single template molecule), and 3) the complexity of the library that is eventually sequenced, in frequencies of clone molecules (how many of each set of clones is selected by the sequencer). The PCR amplification model that results in the amplified library complexity (#2 above) is implemented based on the inherited efficiency model defined in (Best, Oakes, Heather, Shawe-Taylor, & Chain, 2015). The distribution defines the number of times the pool of loci is sampled, the number of duplicate molecules that are generated from a RAD locus template, and the distribution of PCR errors in the resulting reads. The user can define the number of PCR cycles present in the simulation run (by default, no PCR duplicates are generated). The key parameters that define library complexity, including the ratio between pre-amplification DNA template molecules to sequenced reads, can also be specified in the available advanced options. For the *in silico* sequencing of each sample, *RADinitio* iterates over the size of the sequenced library distribution generated by the previously described PCR model. In each iteration, a sequence from the RAD allele pool is picked at random, and a PCR duplicate/error value pair is sampled from the distribution.

For sdRAD libraries, where molecule size is determined by mechanical shearing, an insert size is drawn from a normal distribution for each pair of reads, and a copy of the selected sequence is truncated to the determined length (Fig. 2E). For ddRAD, the inserts are determined by the restriction sites of the corresponding source locus (Fig. 2F). The inserts are then split into a sequenced pair of reads (by default, 2×150bp). The sequence is copied multiple times, to reflect its number of PCR clones, with shared PCR errors added accordingly. Random sequencing errors are finally added to each individual read pair based on an error distribution that follows the normal error pattern observed in Illumina sequencing, where the 3’ end of the read accumulates more errors than the 5’ end (Glenn, 2011; Minoche, Dohm, & Himmelbauer, 2011; Schirmer et al., 2015). The model linearly increases the frequency of errors from the 5’ frequency to the 3’ value through the length of the read. Individual read IDs contain information about the original RAD locus and allele it originated from, its insert size, clone identifier, and duplicate number. The results of the pipeline are paired-end, per-sample FASTA files.

### Comparing RADinitio against empirical RAD datasets

*RADinitio* simulated libraries were retrospectively modeled after empirical RAD datasets to compare the read distribution and coverage obtained from simulated RAD loci. We selected two empirical RAD datasets that represent distinct applications of RAD protocols in the same biological system, the threespine stickleback, *Gasterosteus aculeatus*. First, we modeled a RAD dataset from (Nelson & Cresko, 2018), which contains a single digest, *PstI* library, with 2×250bp paired-end reads and a narrow insert size (380bp±30bp), which was designed for high overlap of forward and reverse reads. Second, we modeled a double digest library made with *NlaIII* and *MluCI*, an insert size range between 295bp and 340bp, and sequenced using 2×100bp paired-end reads from (Stuart et al., 2017).

Reads from both empirical and simulated datasets were aligned to the threespine stickleback reference genome (BROADS1, Ensmbl version 92) using *BWA-MEM* (Li, 2013; Li & Durbin, 2009), converted to BAM files and sorted using *samtools* (Li et al., 2009), and assembled as RAD loci using *Stacks* v2.4 (Rochette et al., 2019). We filtered alignments by removing RAD loci present in less than 80% of all sequenced individuals using the *Stacks* `populations` module. The insert size and coverage distribution for all datasets was obtained using the detailed output of the *Stacks* `gstacks` module.

### Assessing number of RAD loci across reference genomes

We calculated the number of RAD loci obtained under different library configurations across different reference genomes. We selected species with previously published RAD datasets: the threespine stickleback *Gasterosteus aculeatus* (Bassham et al., 2018; Hohenlohe et al., 2010; Nelson & Cresko, 2018; Stuart et al., 2017) the yellow warbler *Setophaga petechia* (Bay et al., 2018), the postman butterfly *Heliconius melpomene* (Davey et al., 2017; Nadeau et al., 2014), and the bush monkeyflower *Mimulus aurantiacus* (Chase, Stankowski, & Streisfeld, 2017; Stankowski et al., 2019). We used available reference sequences for each of the species: stickleback – BROADS1 (Ensembl version 92), *Heliconius* – Hmel2.5 (Lepbase version 4), warbler – draft assembly (Bay et al., 2018), monkeyflower – chromosome-level assembly (Stankowski et al., 2019). When running *RADinitio*, we only simulated over chromosome sequences for stickleback, *Heliconius*, and monkeyflower; only for scaffolds larger than 5Kb for the yellow warbler genome. The --`tally`-`rad`-`loci` command from *RADinitio* was used, which provides the number of RAD loci in a reference sequence under defined library parameters. For all genomes, we simulated sdRAD libraries using the GC-rich 8-bp cutter *SbfI* (CC/TGCAGG), the GC-rich 6-bp cutter *PstI* (C/TGCAG), and the AT-rich 6-bp cutter *EcoRI* (G/AATTC). ddRAD libraries were also generated for each of the main forward cutters, using both the AT-rich 4-bp *MseI* (T/TAA) and the GC-rich 4-bp *MspI* (C/CGG) reverse cutters. For each two-enzyme combination, ddRAD libraries were simulated using a narrow (75bp), intermediate (150bp), and broad (300bp) insert size range, all centered around 350bp.

### Estimating RAD loci number in the absence of a reference

To mimic the process of estimating the number of expected loci in a species with no reference genome, we compared the number of empirical *SbfI* RAD loci in the brown trout *Salmo trutta*, as reported in the dataset published by (Paris et al., 2017), against close salmonids and other related Teleost fishes. Genomes for the Atlantic salmon *Salmo salar* (ICSASG_v2), rainbow trout *Oncorhynchus mykiss* (Omyk_1.0), grayling *Thymallus thymallus* (ASM434828v1), arctic char *Salvelinus alpinus* (ASM291031v2), and Northern Pike *Esox lucius* (Eluc_V3) were downloaded from NCBI. For these genomes, we filtered non-chromosomal scaffolds and estimated *SbfI* RAD loci using *RADinitio* --`tally`-`rad`-`loci`. In addition, an estimate of the number of loci based on the predicted frequencies of *SbfI* sites in the genome – calculated estimating the frequency of a 8-bp long sequence in each of the genomes, with and without taking into account GC content – was generated for all genomes using custom python scripts.

### Allele dropout estimation under different sequencing parameters

We performed two different simulations over the stickleback genome to generate populations with different base levels of genetic polymorphism using the *RADinitio* --`make`-`population` command. The first run simulated four subpopulations, each with an effective population size of 5,000, a mutation rate of 7×10^−8^ mutations per base per generation, a recombination rate of 3×10^−8^ recombination events per base per generation, a symmetric per-generation migration rate of 0.001, and an indel probability of 0.01. From each population, we sampled 25 diploid individuals, for a total of 100 sequenced samples. The second run simulated a population under the same parameters, except for an effective population size of 25,000 to increase genetic polymorphism. The expected genome-wide nucleotide diversity (*π*) in these populations was 0.0058 and 0.0114, respectively.

For these two populations, we simulated both single- and double-digest RAD libraries under different DNA template and sequencing conditions. sdRAD libraries were generated using the *SbfI* restriction enzyme, a mean insert size of 350bp (±35bp), and 2×150 paired-end reads. The first library was generated *in silico* using ‘good’ quality source DNA (defined as having a large number of DNA templates, or a template to sequence ratio of 4:1), PCR amplified for 12 cycles, and sequenced at 15x depth of coverage. The second library had ‘poor’ starting DNA (defined as a small number of DNA templates, or a template to sequence ratio of 1:4), was amplified for 12 PCR cycles and sequenced at 15x depth. The third library had poor-quality starting DNA, was amplified for 12 PCR cycles and sequenced at 30x depth. Quality of DNA is represented by the library diversity in the form of the ratio of DNA template molecules to sequenced DNA reads. The template to sequence ratio broadly represents the quality of the input-DNA, where a small ratio represents a poor template pool and results in a high number of clone (duplicate) molecules after enrichment and sequencing. This process was repeated for ddRAD libraries using the *SbfI-MspI* enzyme combination, using the same insert, DNA quality, and enrichment parameters as described for the sdRAD libraries.

Across the 12 simulated libraries we quantified allele dropout at two different stages of the library. The first stage occurs at the individual allele level, and represents alleles dropped due to genetic polymorphisms in the population – the most common example are alleles that are naturally absent due to the presence of mutations in restriction enzyme cutsites. *RADinitio* provides a tally of allelic dropout due to the underlying genetic variation. The second stage occurs when alleles are lost due to the random sampling associated with library enrichment and sequencing. Variation in DNA quality, enrichment, and sequencing coverage introduce biases in the otherwise even sampling, potentially leading to dropped sequences. Since all the simulated reads contain meta information regarding their allele of origin, we can identify the status of an allele across samples. For our analyses, any allele with a final coverage of 2 or less (after removing PCR duplicates) is considered as dropped, as they will be effectively invisible to any downstream analysis. For both stages of allelic dropout, we quantified the distribution of the percentage of individuals recovered per allele.

## Results

### RADinitio simulations resemble empirical RAD loci

To confirm that the loci reconstructed from *RADinitio*-simulated reads behave similarly to empirical RAD loci, we performed simulations modelled after two published threespine stickleback RAD datasets (Nelson & Cresko, 2018; Stuart et al., 2017) and compared the distribution of recovered insert sizes and average sequencing coverage across loci. We performed two simulations, both representing applications of distinct RAD protocols and sequencing strategies. The first, a single-digest *SbfI* RAD library with 2×250bp paired reads and narrow, highly overlapping insert size distribution from (Nelson & Cresko, 2018). The second, a double-digest library generated with *NlaIII* and *MluCI*, with an insert size range between 295bp and 340bp, and sequenced using 2×100bp paired-end reads from (Stuart et al., 2017).

Reads from the empirical sdRAD library are generated from molecules generated from a narrow insert size (Fig. 3A – red line). Due to the long read lengths of this dataset, the majority of paired-end reads overlap, generating a continuous distribution of coverage between the single- and paired-end regions of the locus (Fig. 3A – blue polygons). Our simulated reads generated under similar conditions *in silico* (Fig. 3C) replicate the insert and coverage distributions of the empirical data. By sampling insert sizes from a normal distribution, *RADinitio* replicates the random shearing and size selections of molecules in the library preparation, which leads to a bell-shaped distribution of both insert size and coverage across the paired-end region of the locus. This sampling process replicates the variation in inserts between loci and across individuals successfully mimicking mechanical shearing in real single-digest libraries (Fig. S2).

**Figure 3.**
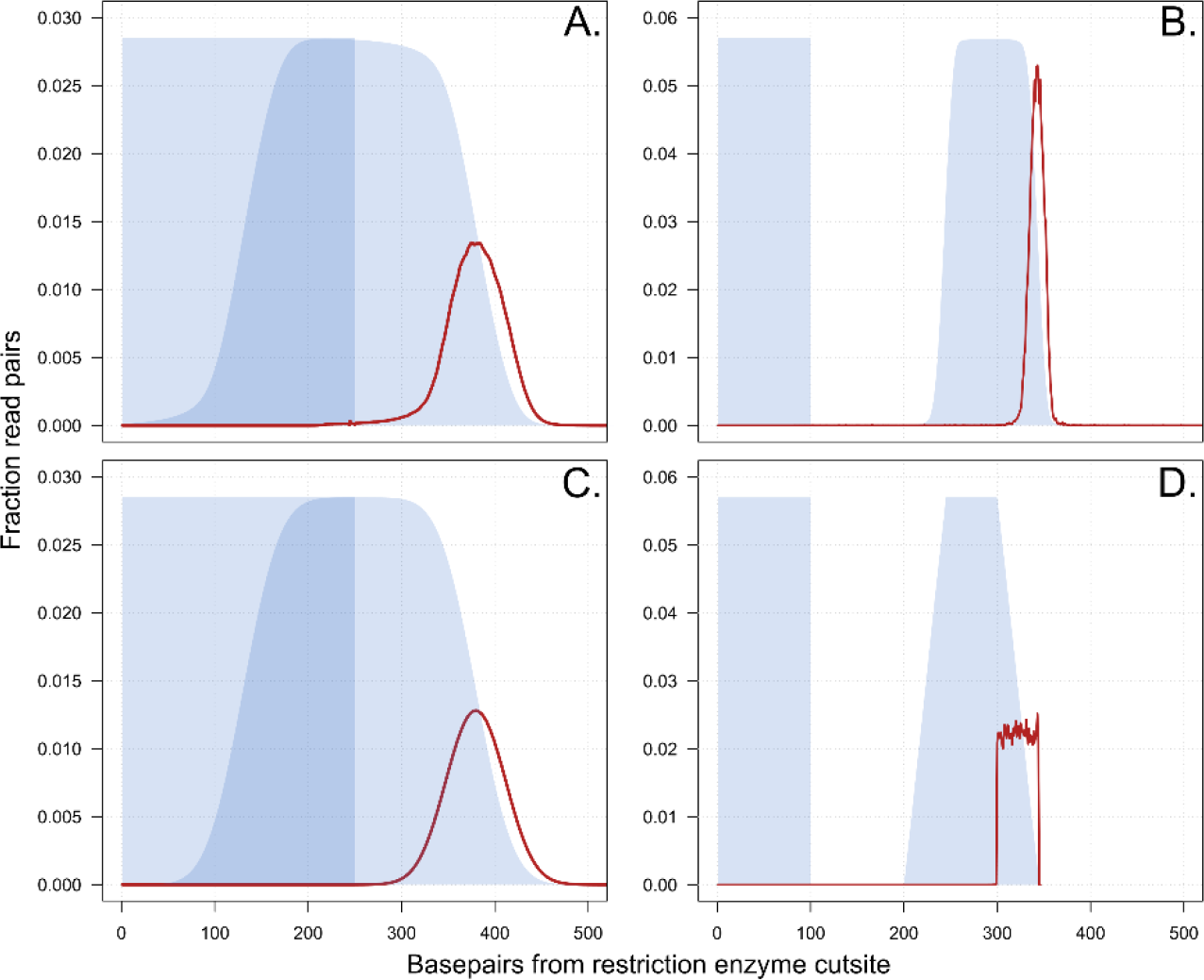
RADinitio simulations are comparable to empirical RAD datasets. Insert size and coverage distributions for two empirical RAD datasets – the sdRAD *PstI* 2×250bp library from Nelson & Cresko (2018) (A), and the ddRAD *NlaIII*-*MluCI* 2×100bp library from Stuart et al. (2017) (B), with their corresponding simulated libraries (C and D), respectively. Red line indicates insert size distributions in the libraries, as determined by assembly of the RAD loci. Blue polygons show sequencing coverage distribution across the single and paired reads in the locus (left and right polygons, respectively), calculated by placing the first base of the restriction site at position 1 and accounting for the dataset’s read length.

In contrast, the 100bp ddRAD reads generate a coverage distribution in which the 5’ and 3’ ends of any given locus are sequenced, but no overlap occurs between single- and paired-end reads (Fig. 3B,D – blue polygons). Variation in inserts and coverage across the paired-end region of the locus reflects both variation in insert sizes across loci, but also variation in single loci if more than one restriction cutsite is retained in the insert range (Fig. S3). In the empirical ddRAD data, we observed a high variation of insert sizes across samples due to what appears to be spurious, non-restriction enzyme-related reads (not shown). As we filtered spurious alignments and kept loci retained in 80% of samples, the empirical insert distribution has a slightly narrower mean insert length relative to the data generated by RADinitio (Fig. 3B – red line).

*RADinitio* can accurately model the behavior of RAD loci across different library preparations and sequencing strategies. Simulating different library and sequencing strategies – altering protocol of choice, insert size distribution, and read lengths – can help guide the preparation of real libraries.

### RADinitio can quantify number of RAD loci across different reference genomes

When designing a RADseq-based experiment, it is essential to know the extent of a genome a RAD library will cover, based on the number of loci present, so that the proper amount of required sequencing can be determined. When a reference genome is available, this estimation is straightforward as it mostly requires quantifying the number of restriction enzyme recognition sequences in the genome of interest. Aside from counting the number of restriction sites present in the genome, *RADinitio* takes a locus-based approach in which the two tags generated from a single restriction site (one on each strand of DNA) are treated as independent objects that can interact with other elements in the library simulation process, such as proximity to other tags and their relative position in a given insert distribution. This allows for detailed estimation of the number of RAD loci in insert size-dependent libraries, such as ddRAD, or to model the behavior of loci when restriction sites are frequent and close to one another.

Across the different species studied, the overall number of RAD loci is roughly proportional to the size of the genome. The yellow warbler assembly (1.26Gb in length) has an overall higher number of loci than the other species (Fig. 4C). Within a single species, the number of loci can vary by an order of magnitude, depending on the enzyme employed. In the threespine stickleback (Fig. 4A), we recover 35,871 and 265,048 *SbfI* and *PstI* loci, respectively. Similar results are observed in *Heliconius*, with 1,942 *SbfI* and 136,011 *EcoRI* loci. Interestingly, while we expect the GC content of the genome to play a role in the enzyme restriction site frequency, we observed that for both stickleback (Fig. 4A) and warbler (Fig. 4C), the number of *PstI* loci (6bp, GC-rich cutter), was higher than *EcoRI* loci (6bp AT-rich cutter), even when both genomes are around 40% GC.

**Figure 4.**
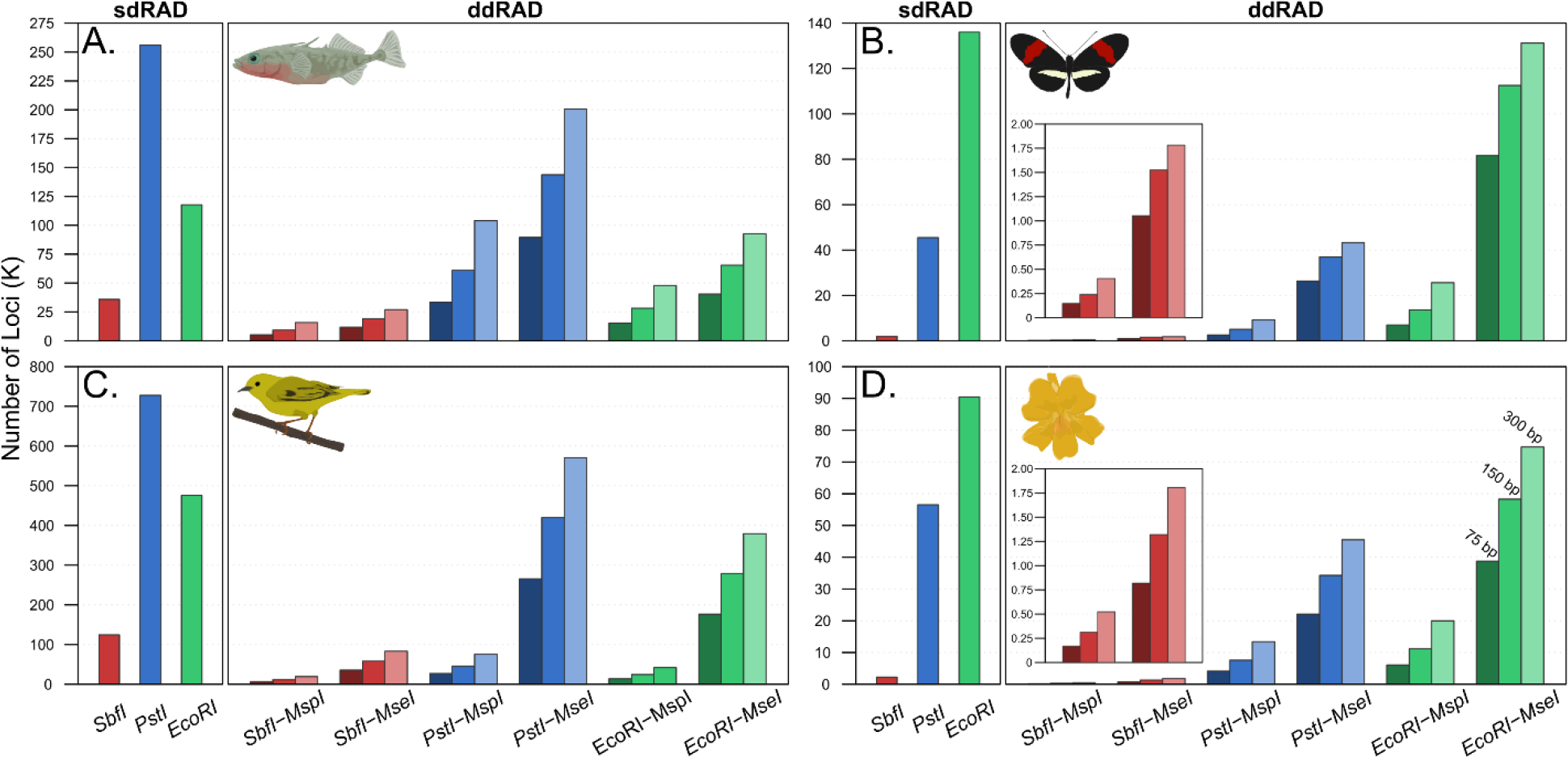
RADinitio can estimate the number of RAD loci recovered from a specific library preparation across reference genomes. RAD libraries were simulated in the genomes of the threespine stickleback (A), *Heliconius melpomene* butterflies (B), yellow warbler (C), and bush monkeyflower (D). Red, blue, and green bars represent libraries with *SbfI, PstI*, and *EcoRI* as primary cutters, respectively. For ddRAD, the restriction enzymes *MspI* and *MseI* are used in combination with each of the main cutters. Dark, intermediate, and light-colored bars represent 75bp, 150bp, and 300bp insert sizes windows, respectively. Inserts in (B) and (D) show number of RAD loci in *SbfI-MspI* and *SbfI-MseI* libraries.

For ddRAD libraries, the number of loci recovered was proportional to the frequency of the rare cutter in the genome. *SbfI* sites are rare across both *Heliconius* (Fig. 4B) and *Mimulus* (Fig. 4D), and therefore ddRAD loci with *SbfI* as the main cutter are even less frequent. Across all genomes, loci were more frequent for *MseI*, an AT-rich cutter, which follows the expectation given the GC frequency of the studied genomes. Across all simulated ddRAD libraries the number of recovered loci is proportional to the library insert size, which is expected given that larger windows provide more sequence within which the secondary restriction site can occur.

### Modelling RAD loci in the absence of a reference genome

Given the suitability of RADseq protocols in non-model systems, many RAD analyses are performed on species lacking a reference genome. If a reference is not available, *RADinitio* simulations can be applied to closely related species in order to explore the expected behavior of the empirical data. A RAD analysis in the brown trout (*Salmo trutta*) (Paris et al., 2017) recovered between 60-85,000 *SbfI* RAD loci, depending on the filtering parameters applied to the data (Fig. 5). Lacking a brown trout reference genome, Paris and colleagues used information about the number of RAD loci observed in the Atlantic salmon, a congener to the brown trout, to optimize *de novo* assembly parameters post-data generation.

**Figure 5.**
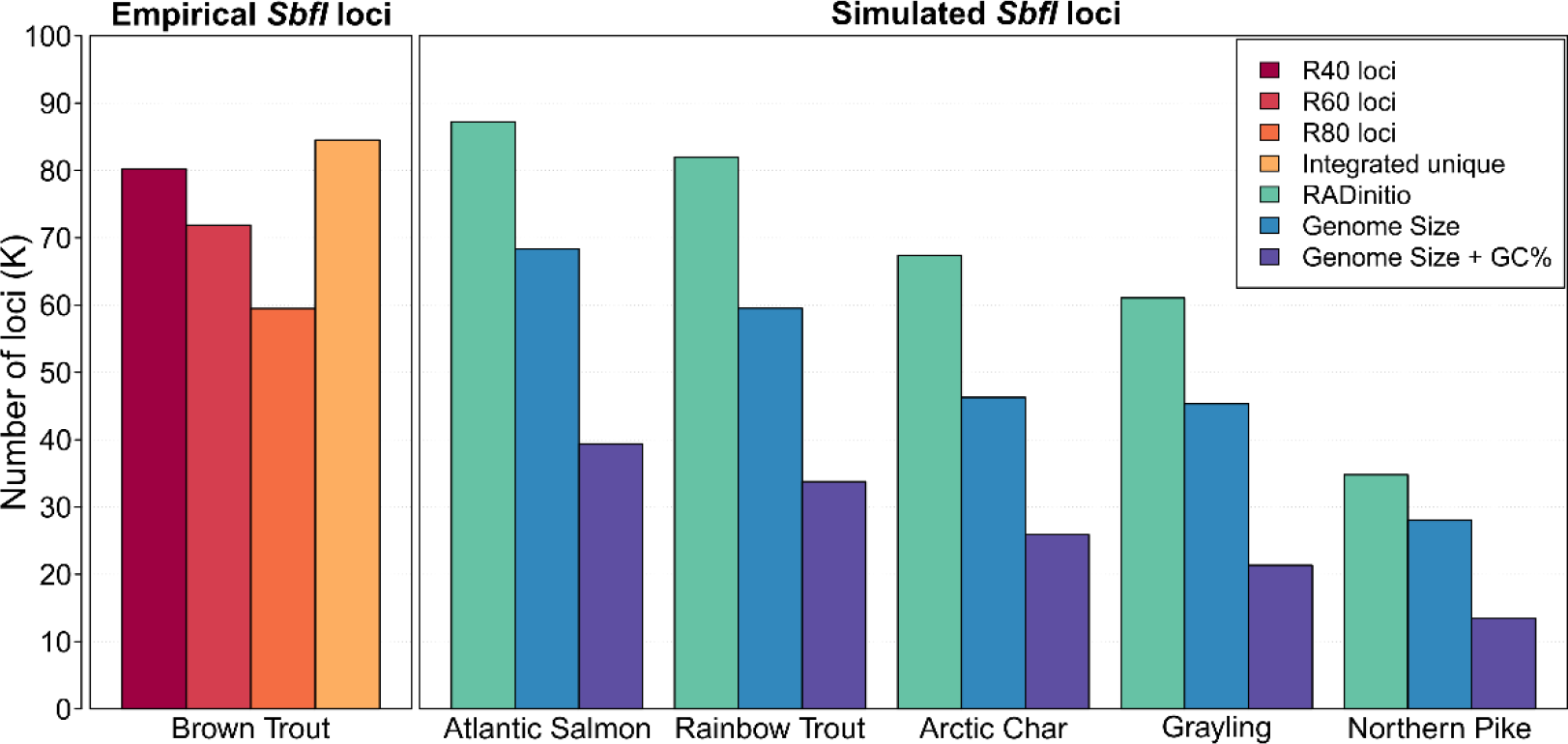
RADinitio can estimate the number of RAD loci de novo. Left panel, empirical *SbfI* RAD loci recovered in the Brown Trout by Paris et al. (2017). Colors of the bars represent the metric used in their analysis to retain loci. R40, R60, and R80 represent loci present in 40%, 60%, and 80% of individuals, respectively. Integrated unique loci are loci recovered in the brown trout that mapped uniquely to the Atlantic salmon genome. Right panel, number of recovered *SbfI* RAD loci from simulations across several related species. Green bars represent the number of loci recovered with *RADinitio*. Blue and purple bars represent the number of loci found based on the frequency of the *SbfI* recognition sequence in the species’ genome using both 50% and the genome’s actual GC content (40-42%), respectively.

A similar simulation framework can be applied to estimate properties of the data before sequencing, most commonly quantifying the number of expected markers to account for limitations in sequencing. Expanding the framework used by (Paris et al., 2017), we quantified the expected number of *SbfI* RAD loci across species closely related to the brown trout and for which a genome is available (Fig. 5), using various estimation parameters. First, species estimation methods that only considered the size of the genome and nucleotide frequencies underestimate the actual number of restriction sites. For example, across the Atlantic salmon chromosomes *RADinitio* identifies 87,198 *SbfI* RAD loci, while estimating just using genome size and genome size plus GC% returns between 68,366 and 39,392 loci, respectively. When we compare our estimates against the empirical data, we observe that across all species, estimates based on the count of *SbfI* sites are of similar magnitude to empirically recovered loci across various levels of missing data filtering in the empirical brown trout dataset (Fig. 5).

While the overall similarity in locus number follows evolutionary relatedness (Lien et al., 2016), given genome evolution in Salmonids, some patterns are more complex. As expected, the Atlantic salmon displays the most similar *SbfI* RAD loci distribution to the observed brown trout empirical data. The rainbow trout and Arctic char are both in the sister clade to *Salmo* but display different *SbfI* loci numbers. Under this metric, the char’s locus distribution appears more similar to the grayling’s despite greater evolutionary distance. They do, however, share similar rediploidization histories following the salmonid genome duplication (Christensen et al., 2018; Varadharajan et al., 2018) and comparable genome architecture. The northern pike, a species sister to salmonids, has a smaller number of loci, which is expected given its smaller genome, and would thus not serve as an adequate model for estimating RAD loci in the brown trout unless there were no alternatives. By performing estimates based on a variety of related species of different evolutionary distances, we can bound our expectations for our genome of interest.

### Effects of polymorphism and library preparation on allele dropout

The effects of allelic dropout have been extensively discussed in the literature as one of the main caveats in RADseq (Andrews et al., 2016; Arnold, Corbett-Detig, Hartl, & Bomblies, 2013; Davey et al., 2013; Puritz, Matz, et al., 2014). Mutations to a restriction site sequence can cause variation in the presence or absence of a particular RAD locus within and among populations and individuals (Arnold et al., 2013). Depending on the scale, this allelic dropout can affect the estimation of population genetic parameters (Arnold et al., 2013). However, the prevalence and impact of allelic dropout due to mutations to a restriction site have not been extensively studied. Moreover, how much contribution genetic polymorphism has over dropout, in comparisons to components of library preparation and sequencing, is not yet fully understood. To test the effects of genetic polymorphism and library preparation and sequencing on allele dropout, we simulated populations with low (Fig. 6 – columns 1 & 3) and high polymorphism (Fig. 6 – columns 2 & 4), across sdRAD (Fig. 6 – columns 1 & 2) and ddRAD libraries (Fig. 6 – columns 3 & 4).

**Figure 6.**
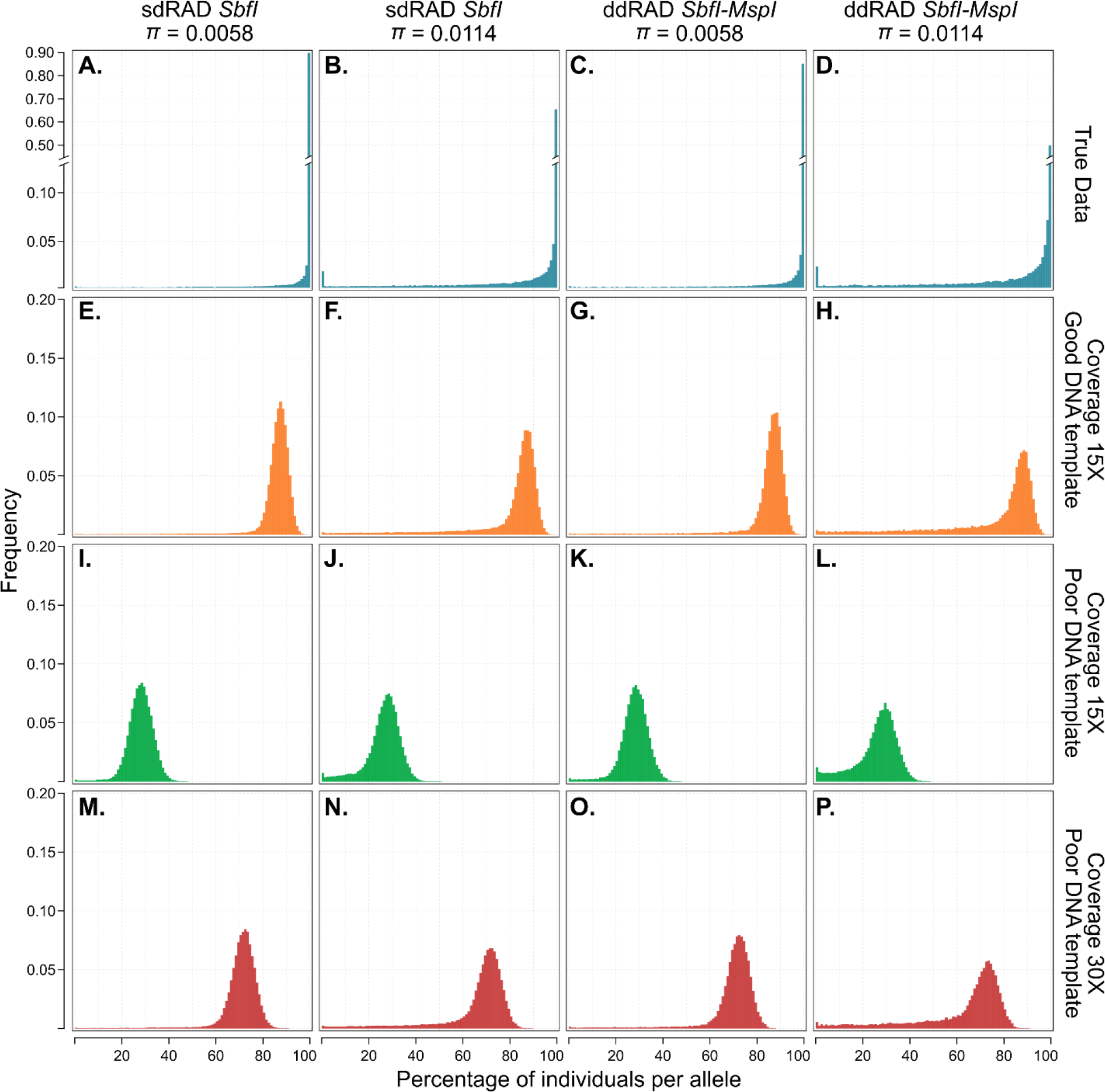
Effects of genetic polymorphism, RAD library preparation, and sequencing on allelic dropout. Histograms display the distribution of the percentage of individuals per allele – a function of library-wide missing data – across experimental conditions. Each column shows a different configuration of genetic diversity and RAD library protocol. Top row represents the true state of the data – the base level of missing data based just on polymorphism-based allelic dropout. Rows 2 to 4 show subsequent levels of missing data due to different combinations of sequencing depth and DNA quality, which represent library preparation and sequencing-dependent allelic dropout.

When quantifying the presence of individual alleles (Fig. 6 – row 1), ddRAD has an overall lower percentage of samples per allele across both populations (Fig. 6A – 97.7% vs Fig. 6C – 96.8%, and Fig. 6B – 90.1% vs Fig. 6D – 86.0%), representing more allele dropout. Differences are negligible but expected – in sdRAD with *SbfI* there are eight sites for a mutation causing dropout, versus a total of 12 in ddRAD – eight in *SbfI* and four in *MspI*. Additionally, higher chances of dropout are observed in populations with higher polymorphism (Fig. 6A – 97.7% vs Fig. 5B – 90.1%). This is also expected, as a higher polymorphism rate increases the chance of a mutation at any given site, among them restriction enzyme recognition sequences. What is interesting however, is that from an analysis perspective the observed recovery of alleles is above some of the most commonly used filtering thresholds for missing data in population genomic studies. One example, the R80 metric from Paris et al. (2017) filters sequences for which less than 80% of samples are present. In this example, even the lowest observed value for allele-level dropout (Fig. 6D) means that on average an allele is present in 86% of samples, and missing data due only to cut site mutations should have minimal impacts on downstream biological inferences.

The process of library preparation and sequencing behaves as a sampling process for the alleles present in an individual. The effects of library preparation and sequencing introduce biases to this sampling, impacting the alleles that can be recovered downstream. We modeled this bias in sampling by preparing *in silico* RAD libraries enriched and sequenced under variable conditions, originating from the simulated populations described above.

First, we simulated a library based on good quality DNA (see Methods for a definition of ‘good’ and ‘poor’ quality) and sequenced at a coverage of 15x (Fig. 6 – row 2). Coverage of 15x means that, on average, an allele would be sequenced 7.5 times if no other biases were present. Because DNA quality is good, the effect of PCR duplicates is minimal, but sampling biases due to relatively low per-allele coverage result in an increase in allele dropout of around 10% (for example, Fig. 6A – 97.7% vs Fig. 6E – 85.8%). This alone accounts for increases in dropout greater to those observed due to of increases in polymorphism (Fig. 6A – 97.7% vs. Fig. 6B – 90.1%).

Poor DNA template quality, which can bias library enrichment, increases dropout by up to 60% (Fig.6 – Row 3). While sequencing coverage is still 15x, there are fewer DNA templates, so the sequencing capacity is filled by more PCR-generated duplicates. After analytically removing duplicates (since they add no new information), the depth of each locus falls to 3-5x, which limits the even sampling of alleles and results in dropout even when the allele is present in the individual. In contrast, while DNA quality is still poor, an increase in total sequencing depth to 30x (Fig. 6 – row 4) results in a post-duplicate removal depth of coverage of 8-10x. While still low for many downstream analyses, this resulting coverage allows for the recovery of alleles in 60 to 70% of samples.

This assessment suggests that the sampling properties of the library preparation process extensively contributes to the loss of alleles. While allele dropout due to natural polymorphism is real and has an effect over downstream biological inferences, at a population level, their presence can be indistinguishable from sampling biases produced by library preparation and sequencing.

## Discussion

### Choosing the right RAD protocol matters

The success of a RADseq experiment strongly depends on the selection of a library protocol and restriction enzyme(s), as these define the genetic markers used post-analysis for reaching biological inference. Given the variation in the magnitude of RAD loci obtained across different genomes (Fig. 4) it is important for researchers to choose an enzyme/protocol that suits their biological system and experimental design. Choosing a set of restriction enzymes that cut the genome too infrequently could lead to a reduced sampling of linkage blocks across chromosomes. This reduced sampling has been subject to arduous debate in the RADseq community (Catchen et al., 2017; Lowry et al., 2016; McKinney, Larson, Seeb, & Seeb, 2017) and could be detrimental for certain experimental designs – such as when identifying signatures of selection in a genome. On the opposite side of the spectrum, massively oversampling the genome could lead to problems with reduced sequencing coverage and more allelic dropout when the experimenter is not aware of the magnitude of the data generated per-individual.

Additionally, even after choosing the right RAD protocol and restriction enzyme(s), researchers need to be aware of the effects of other library preparation components – such as insert sizes – on the data to be generated. While insert size selection does not have an effect in the recovery of loci in sdRAD libraries, it can impact assembly and coverage of loci when performing paired-end sequencing (Rochette et al., 2019). Narrow insert ranges will reduce the length of the resulting loci, while broad ones can negatively impact coverage across the paired-end reads generated from random shearing. In contrast, locus recovery is affected by insert sizes in ddRAD libraries – broader inserts allow for more secondary restriction sites to be recovered across more molecules, causing more loci to be kept but leading to increased variability in coverage across paired-reads. The original ddRAD protocol (Peterson et al., 2012) used a narrow insert range between 50 to 100bp, in part because of limitations in sequencing capacity at the time of the publication. Given an increase in capacity in modern Illumina sequencers, the protocol might not be limited to these narrow size ranges in current ddRAD experiments.

Advances in sequencing capacity, as well as increased accessibility to resources for library preparation procedures mean that some of the concepts outlined in these original protocol publications (Baird et al., 2008; Etter et al., 2011; Peterson et al., 2012) might be outdated for the preparation of current RAD libraries. Understandably, researchers will use the protocol that is the most accessible, often using the same library procedures a collaborator has used in the past. While we do not advocate against this practice, our intention is for investigators to explore their protocol of interest on their system of study – and its suitability in their biological system and experimental design – before making the time and monetary investment in library preparation and sequencing. *RADinitio* allows for this exploration of protocol selection, restriction enzyme(s), and sequencing requirements in the early stages of the experimental process.

### How to properly mitigate allelic dropout

The absence of alleles due to the presence of polymorphisms in individuals of the study population is an inevitable aspect of RADseq experiments. However, this fact does not limit the capacity of RADseq as a tool for population and conservation genetics, and phylogenomics. Poor template DNA, as well as errors in molecular library generation and sequencing strategy, are likely to account for a greater portion of missing data than natural polymorphism. Our simulations highlight that most issues encountered by experimenters may be due to problems with DNA quality, library preparation, and sequencing, and not by genetic variation within their biological system. The effects of poor data generation are often indistinguishable from true biological noise and without detailed assessment of the sequencing data – via verifying coverage across samples, removing duplicate sequences, testing filtering parameters, among others – might go unnoticed.

Recovering RADseq alleles is dependent on several sampling processes during library preparation and sequencing. Sequencing depth of coverage impacts this process by determining the total number of molecules sampled. At low coverages, sequences from one allele might be sampled more just by chance, effectively causing dropout in the other allele. DNA quality and quantity can exacerbate these sampling biases, by limiting the number of original molecules available for enrichment and sequencing, effectively limiting the information capacity of the library. When the amounts of template DNA are small, the *genomic molarity* of a sample – the number of haploid genome copies present – is low. Each of these haploid copies of the genome, which contain unique information, physically limit the number of times an allele can be uniquely sampled. Enrichment by PCR, perhaps the most common method used to mitigate the effects of low genomic molarity, further bias this process by masking library richness – making it seem like more starting DNA is available – and/or by differentially amplifying alleles. While increasing sequencing depth will not negate the effects of PCR-based artifacts, if can improve the post-duplicate removal coverage and the recovery of alleles (Fig. 6 – row 4). In situations in which clones far outnumber unique molecules the increased coverage merely allows for a higher number of unique molecules to be sequenced. However, the information content of the library will always be limited by its genomic molarity, and new molecules will not be obtained if they are not available even after increased sequencing coverage.

The estimation of genomic molarity is complicated in RADseq, as only a portion of the genome is informative. The total mass of DNA used in the experiment, while representative of total genomic molarity, will overestimate the number of available RAD loci. For example, while a library can be prepared using 1μg of DNA per sample, this mass accounts for DNA that is anchored by a restriction site (which will be sequenced), and a fraction of DNA that is not. Depending on the density of markers selected, the mass of RAD loci will often be orders of magnitude lower. While some RAD protocols account for this by physical enrichment of RAD loci (Ali et al., 2016), most depend on PCR amplification for their enrichment, leading to the sampling problems described above. The problems with low quality or quantity of starting DNA template are further increased in RAD (or RAD-like) protocols that require small amounts of DNA and rely heavily on extensive PCR amplification (Barilli et al., 2018; Elshire et al., 2011; Kilian et al., 2012). DNA accessibility permitting, researchers should be able to ameliorate the described problems by starting their library preparations with as much starting template as possible – minimizing the reliance on extensive PCR amplification.

In our simulations, we see that problems with DNA quality and sequencing depth can lead to up to a 60% increase in allele dropout, under specific circumstances. The effects in real biological data could be even larger given the stochastic nature of many of the biochemical reactions and random sampling that occurs in the library preparation process, which is impossible for our simulations to truly model. Regardless of the limitations, *RADinitio* provides a tool for the exploration of possible sources of error prior to the preparation and sequencing of RADseq libraries. While software exists for the optimization of data post sequencing (Ilut et al., 2014; McCartney-Melstad et al., 2019; Paris et al., 2017; Rochette & Catchen, 2017), their use is meant for the selection of analytical parameters. This optimization will definitely improve the recovery of data when it is available but cannot – and is not designed to – salvage data when the aforementioned problems are widespread. Prospectively simulating a RADseq experiment can reveal sources of error and elucidate how a particular protocol will interact with an organism of interest. Simulation prior to sequencing can provide a roadmap for a proposed experiment and ensure a sound analytical foundation prior to the investment of resources.

## Acknowledgments

The authors would like to thank José Cerca for coming up with the *RADinitio* name and for his assessment of the Stacks mailing list. Matt Streisfeld for early access to the *M. aurantiacus* reference assembly, and Yoel Stuart and Daniel Bolnick for the stickleback ddRAD data. We also want to thank Niraj Rayamajhi, Jan Stefka, Eric Normandeau for early testing of the software. AGR and NCR were supported by NSF grant 1645087.

## Data Accessibility

*RADinitio* is released under the free software, GPLv3 license. The software and documentation are available from the Catchen Lab website and from the Python Package Index (PyPI).

## Author Contributions

JMC, NCR, and AGRC designed *RADinitio*, designed and revised experiments, and wrote the manuscript. AGRC implemented *RADinitio* along with NCR. AGRC conducted the experiments.

## References

Ali, O. A., O’Rourke, S. M., Amish, S. J., Meek, M. H., Luikart, G., Jeffres, C., & Miller, M. R. (2016). RAD Capture (Rapture): Flexible and Efficient Sequence-Based Genotyping. Genetics, 202(2), 389–400. doi: 10.1534/genetics.115.183665

Amores, A., Catchen, J. M., Ferrara, A., Fontenot, Q., & Postlethwait, J. H. (2011). Genome evolution and meiotic maps by massively parallel DNA sequencing: Spotted gar, an outgroup for the teleost genome duplication. Genetics, 188(4), 799–808. doi: 10.1534/genetics.111.127324

Amores, A., Wilson, C. A., Allard, C. A. H., Detrich, H. W., & Postlethwait, J. H. (2017). Cold Fusion: Massive Karyotype Evolution in the Antarctic Bullhead Notothen Notothenia coriiceps. G3&#58; Genes|Genomes|Genetics, 7(7), 2195–2207. doi: 10.1534/g3.117.040063

Andrews, K. R., Good, J. M., Miller, M. R., Luikart, G., & Hohenlohe, P. A. (2016). Harnessing the power of RADseq for ecological and evolutionary genomics. Nature Reviews. Genetics, 17(2), 81–92. doi: 10.1038/nrg.2015.28

Arnold, B., Corbett-Detig, R. B., Hartl, D., & Bomblies, K. (2013). RADseq underestimates diversity and introduces genealogical biases due to nonrandom haplotype sampling. Molecular Ecology, 22(11), 3179–3190. doi: 10.1111/mec.12276

Baird, N. a., Etter, P. D., Atwood, T. S., Currey, M. C., Shiver, A. L., Lewis, Z. a., … Johnson, E. a. (2008). Rapid SNP Discovery and Genetic Mapping Using Sequenced RAD Markers. PLoS ONE, 3(10), e3376. doi: 10.1371/journal.pone.0003376

Barilli, E., Cobos, M. J., Carrillo, E., Kilian, A., Carling, J., & Rubiales, D. (2018). A High-Density Integrated DArTseq SNP-Based Genetic Map of Pisum fulvum and Identification of QTLs Controlling Rust Resistance. Frontiers in Plant Science, 9. doi: 10.3389/fpls.2018.00167

Bassham, S., Catchen, J., Lescak, E., von Hippel, F. A., & Cresko, W. A. (2018). Repeated Selection of Alternatively Adapted Haplotypes Creates Sweeping Genomic Remodeling in Stickleback. Genetics, genetics.300610.2017. doi: 10.1534/genetics.117.300610

Bay, R. A., Harrigan, R. J., Underwood, V. Le, Gibbs, H. L., Smith, T. B., & Ruegg, K. (2018). Genomic signals of selection predict climate-driven population declines in a migratory bird. Science, 359(6371), 83–86. doi: 10.1126/science.aan4380

Best, K., Oakes, T., Heather, J. M., Shawe-Taylor, J., & Chain, B. (2015). Computational analysis of stochastic heterogeneity in PCR amplification efficiency revealed by single molecule barcoding. Scientific Reports, 5, 14629. Retrieved from http://dx.doi.org/10.1038/srep14629

Campbell, E. O., Brunet, B. M. T., Dupuis, J. R., & Sperling, F. A. H. (2018). Would an RRS by any other name sound as RAD? Methods in Ecology and Evolution, 0(0). doi: 10.1111/2041-210X.13038

Catchen, J. M., Amores, A., Hohenlohe, P. A., Cresko, W. A., Postlethwait, J. H., & De Koning, D.-J. (2011). Stacks: Building and Genotyping Loci De Novo From Short-Read Sequences. G3: Genes, Genomes, Genetics, 1(3), 171–182. doi: 10.1534/g3.111.000240

Catchen, J. M., Hohenlohe, P. A., Bassham, S., Amores, A., & Cresko, W. A. (2013). Stacks: an analysis tool set for population genomics. Molecular Ecology, 22(11), 3124–3140. doi: 10.1111/mec.12354

Catchen, J. M., Hohenlohe, P. A., Bernatchez, L., Funk, W. C., Andrews, K. R., & Allendorf, F. W. (2017). Unbroken: RADseq remains a powerful tool for understanding the genetics of adaptation in natural populations. Molecular Ecology Resources. doi: 10.1111/1755-0998.12669

Chase, M. A., Stankowski, S., & Streisfeld, M. A. (2017). Genomewide variation provides insight into evolutionary relationships in a monkeyflower species complex (Mimulus sect. Diplacus). American Journal of Botany, 104(10), 1510–1521. doi: 10.3732/ajb.1700234

Chiou, K. L., & Bergey, C. M. (2018). Methylation-based enrichment facilitates low-cost, noninvasive genomic scale sequencing of populations from feces. Scientific Reports, 8(1), 1975. doi: 10.1038/s41598-018-20427-9

Chong, Z., Ruan, J., & Wu, C.-I. (2012). Rainbow: an integrated tool for efficient clustering and assembling RAD-seq reads. Bioinformatics, 28(21), 2732–2737. doi: 10.1093/bioinformatics/bts482

Christensen, K. A., Rondeau, E. B., Minkley, D. R., Leong, J. S., Nugent, C. M., Danzmann, R. G., … Koop, B. F. (2018). The Arctic charr (Salvelinus alpinus) genome and transcriptome assembly. PLOS ONE, 13(9), e0204076. doi: 10.1371/journal.pone.0204076

Cristofari, R., Bertorelle, G., Ancel, A., Benazzo, A., Le Maho, Y., Ponganis, P. J., … Trucchi, E. (2016). Full circumpolar migration ensures evolutionary unity in the Emperor penguin. Nature Communications, 7, 11842. Retrieved from http://dx.doi.org/10.1038/ncomms11842

Danecek, P., Auton, A., Abecasis, G., Albers, C. A., Banks, E., DePristo, M. A., … Durbin, R. (2011). The variant call format and VCFtools. Bioinformatics, 27(15), 2156–2158. doi: 10.1093/bioinformatics/btr330

Davey, J. W., Barker, S. L., Rastas, P. M., Pinharanda, A., Martin, S. H., Durbin, R., … Jiggins, C. D. (2017). No evidence for maintenance of a sympatric Heliconius species barrier by chromosomal inversions. Evolution Letters, 1(3), 138–154. doi: 10.1002/evl3.12

Davey, J. W., Cezard, T., Fuentes-Utrilla, P., Eland, C., Gharbi, K., & Blaxter, M. L. (2013). Special features of RAD Sequencing data: implications for genotyping. Molecular Ecology, 22(11), 3151–3164. doi: 10.1111/mec.12084

Dudaniec, R. Y., Yong, C. J., Lancaster, L. T., Svensson, E. I., & Hansson, B. (2018). Signatures of local adaptation along environmental gradients in a range-expanding damselfly (Ischnura elegans). Molecular Ecology, 27(11), 2576–2593. doi: 10.1111/mec.14709

Eaton, D. A. R. (2014). PyRAD: assembly of de novo RADseq loci for phylogenetic analyses. Bioinformatics, 30(13), 1844–1849. doi: 10.1093/bioinformatics/btu121

Eaton, D. A. R., Spriggs, E. L., Park, B., & Donoghue, M. J. (2016). Misconceptions on Missing Data in RAD-seq Phylogenetics with a Deep-scale Example from Flowering Plants. Systematic Biology, syw092. doi: 10.1093/sysbio/syw092

Elshire, R. J., Glaubitz, J. C., Sun, Q., Poland, J. A., Kawamoto, K., Buckler, E. S., & Mitchell, S. E. (2011). A Robust, Simple Genotyping-by-Sequencing (GBS) Approach for High Diversity Species. PLoS ONE, 6(5), e19379. doi: 10.1371/journal.pone.0019379

Etter, P. D., Bassham, S. L., Hohenlohe, P. A., Johnson, E. a., & Cresko, W. A. (2011). SNP Discovery and Genotyping for Evolutionary Genetics Using RAD Sequencing. Methods Mol Biol, 772(6), 157–178. doi: 10.1007/978-1-61779-228-1

Glenn, T. C. (2011). Field guide to next-generation DNA sequencers. Molecular Ecology Resources, 11(5), 759–769. doi: 10.1111/j.1755-0998.2011.03024.x

Haller, B. C., Galloway, J., Kelleher, J., Messer, P. W., & Ralph, P. L. (2019). Tree-sequence recording in SLiM opens new horizons for forward-time simulation of whole genomes. Molecular Ecology Resources, 19(2), 552–566. doi: 10.1111/1755-0998.12968

Haller, B. C., & Messer, P. W. (2019). SLiM 3: Forward Genetic Simulations Beyond the Wright–Fisher Model. Molecular Biology and Evolution, 36(3), 632–637. doi: 10.1093/molbev/msy228

Hoffberg, S. L., Kieran, T. J., Catchen, J. M., Devault, A., Faircloth, B. C., Mauricio, R., & Glenn, T. C. (2016). RADcap: sequence capture of dual-digest RADseq libraries with identifiable duplicates and reduced missing data. Molecular Ecology Resources, 16(5), 1264–1278. doi: 10.1111/1755-0998.12566

Hohenlohe, P. A., Bassham, S. L., Etter, P. D., Stiffler, N., Johnson, E. a., & Cresko, W. a. (2010). Population Genomics of Parallel Adaptation in Threespine Stickleback using Sequenced RAD Tags. PLoS Genetics, 6(2), e1000862. doi: 10.1371/journal.pgen.1000862

Ilut, D. C., Nydam, M. L., & Hare, M. P. (2014). Defining Loci in Restriction-Based Reduced Representation Genomic Data from Nonmodel Species: Sources of Bias and Diagnostics for Optimal Clustering. BioMed Research International, 2014, 1–9. doi: 10.1155/2014/675158

Jeffery, N. W., DiBacco, C., Van Wyngaarden, M., Hamilton, L. C., Stanley, R. R. E., Bernier, R., … Bradbury, I. R. (2017). RAD sequencing reveals genomewide divergence between independent invasions of the European green crab (Carcinus maenas) in the Northwest Atlantic. Ecology and Evolution, 7(8), 2513–2524. doi: 10.1002/ece3.2872

Kelleher, J., Etheridge, A. M., & McVean, G. (2016). Efficient Coalescent Simulation and Genealogical Analysis for Large Sample Sizes. PLOS Computational Biology, 12(5), e1004842. doi: 10.1371/journal.pcbi.1004842

Kelly, T. J., & Smith, H. O. (1970). A restriction enzyme from Hemophilus influenzae. Journal of Molecular Biology, 51(2), 393–409. doi: 10.1016/0022-2836(70)90150-6

Kilian, A., Wenzl, P., Huttner, E., Carling, J., Xia, L., Blois, H., … Uszynski, G. (2012). Diversity Arrays Technology: A Generic Genome Profiling Technology on Open Platforms. In F. Pompanon & A. Bonin (Eds.), Data Production and Analysis in Population Genomics (pp. 67–89). doi: 10.1007/978-1-61779-870-2_5

Lepais, O., & Weir, J. T. (2014). SimRAD: an R package for simulation-based prediction of the number of loci expected in RADseq and similar genotyping by sequencing approaches. Molecular Ecology Resources, 14(6), 1314–1321. doi: 10.1111/1755-0998.12273

Li, H. (2013, March 16). Aligning sequence reads, clone sequences and assembly contigs with BWA-MEM. Retrieved from http://arxiv.org/abs/1303.3997

Li, H., & Durbin, R. M. (2009). Fast and accurate short read alignment with Burrows-Wheeler transform. Bioinformatics, 25(14), 1754–1760. doi: 10.1093/bioinformatics/btp324

Li, H., Handsaker, B., Wysoker, A., Fennell, T. J., Ruan, J., Homer, N., … Durbin, R. M. (2009). The Sequence Alignment/Map format and SAMtools. Bioinformatics, 25(16), 2078–2079. doi: 10.1093/bioinformatics/btp352

Lien, S., Koop, B. F., Sandve, S. R., Miller, J. R., Kent, M. P., Nome, T., … Davidson, W. S. (2016). The Atlantic salmon genome provides insights into rediploidization. Nature, 533(7602), 200–205. doi: 10.1038/nature17164

Lowry, D. B., Hoban, S., Kelley, J. L., Lotterhos, K. E., Reed, L. K., Antolin, M. F., & Storfer, A. (2016). Breaking RAD: An evaluation of the utility of restriction site associated DNA sequencing for genome scans of adaptation. Molecular Ecology Resources. doi: 10.1111/1755-0998.12596

McCartney-Melstad, E., Gidiş, M., & Shaffer, H. B. (2019). An empirical pipeline for choosing the optimal clustering threshold in RADseq studies. Molecular Ecology Resources, 1755–0998.13029. doi: 10.1111/1755-0998.13029

McKinney, G. J., Larson, W. A., Seeb, L. W., & Seeb, J. E. (2017). RADseq provides unprecedented insights into molecular ecology and evolutionary genetics: comment on Breaking RAD by Lowry et al. (2016). Molecular Ecology Resources, 17(3), 356–361. doi: 10.1111/1755-0998.12649

Melo, A. T. O., & Hale, I. (2019). Expanded functionality, increased accuracy, and enhanced speed in the de novo genotyping-by-sequencing pipeline GBS-SNP-CROP. Bioinformatics, 35(10), 1783–1785. doi: 10.1093/bioinformatics/bty873

Minoche, A. E., Dohm, J. C., & Himmelbauer, H. (2011). Evaluation of genomic high-throughput sequencing data generated on Illumina HiSeq and Genome Analyzer systems. Genome Biology, 12(11), R112. doi: 10.1186/gb-2011-12-11-r112

Mora-Márquez, F., García-Olivares, V., Emerson, B. C., & López de Heredia, U. (2017). ddradseqtools: a software package for in silico simulation and testing of double-digest RADseq experiments. Molecular Ecology Resources, 17(2), 230–246. doi: 10.1111/1755-0998.12550

Nadeau, N. J., Ruiz, M., Salazar, P., Counterman, B. A., Medina, J. A., Ortiz-Zuazaga, H., … Papa, R. (2014). Population genomics of parallel hybrid zones in the mimetic butterflies, H. melpomene and H. erato. Genome Research, 24(8), 1316–1333. doi: 10.1101/gr.169292.113

Narum, S. R., Buerkle, C. A., Davey, J. W., Miller, M. R., & Hohenlohe, P. A. (2013). Genotyping-by-sequencing in ecological and conservation genomics. Molecular Ecology, 22(11), 2841–2847. doi: 10.1111/mec.12350

Nelson, T. C., & Cresko, W. A. (2018). Ancient genomic variation underlies repeated ecological adaptation in young stickleback populations. Evolution Letters. doi: 10.1002/evl3.37

Paris, J. R., Stevens, J. R., & Catchen, J. M. (2017). Lost in parameter space: a road map for stacks. Methods in Ecology and Evolution, 8(10), 1360–1373. doi: 10.1111/2041-210X.12775

Peterson, B. K., Weber, J. N., Kay, E. H., Fisher, H. S., & Hoekstra, H. E. (2012). Double Digest RADseq: An Inexpensive Method for De Novo SNP Discovery and Genotyping in Model and Non-Model Species. PLoS ONE, 7(5), e37135. doi: 10.1371/journal.pone.0037135

Portnoy, D. S., Puritz, J. B., Hollenbeck, C. M., Gelsleichter, J., Chapman, D., & Gold, J. R. (2015). Selection and sex-biased dispersal in a coastal shark: the influence of philopatry on adaptive variation. Molecular Ecology, 24(23), 5877–5885. doi: 10.1111/mec.13441

Puritz, J. B., Hollenbeck, C. M., & Gold, J. R. (2014). dDocent : a RADseq, variant-calling pipeline designed for population genomics of non-model organisms. PeerJ, 2, e431. doi: 10.7717/peerj.431

Puritz, J. B., Matz, M. V, Toonen, R. J., Weber, J. N., Bolnick, D. I., & Bird, C. E. (2014). Demystifying the RAD fad. Molecular Ecology, 23(24), 5937–5942. doi: 10.1111/mec.12965

Razkin, O., Sonet, G., Breugelmans, K., Madeira, M. J., Gómez-Moliner, B. J., & Backeljau, T. (2016). Species limits, interspecific hybridization and phylogeny in the cryptic land snail complex Pyramidula : The power of RADseq data. Molecular Phylogenetics and Evolution, 101, 267–278. doi: 10.1016/j.ympev.2016.05.002

Rochette, N. C., & Catchen, J. M. (2017). Deriving genotypes from RAD-seq short-read data using Stacks. Nature Protocols, 12(12), 2640–2659. doi: 10.1038/nprot.2017.123

Rochette, N. C., Rivera-Colón, A. G., & Catchen, J. M. (2019). Stacks 2: Analytical Methods for Paired-end Sequencing Improve RADseq-based Population Genomics. BioRxiv, 615385. doi: 10.1101/615385

Schirmer, M., Ijaz, U. Z., D’Amore, R., Hall, N., Sloan, W. T., & Quince, C. (2015). Insight into biases and sequencing errors for amplicon sequencing with the Illumina MiSeq platform. Nucleic Acids Research, 43(6), e37–e37. doi: 10.1093/nar/gku1341

Schlötterer, C. (2004). The evolution of molecular markers — just a matter of fashion? Nature Reviews Genetics, 5(1), 63–69. doi: 10.1038/nrg1249

Small, C. M., Bassham, S., Catchen, J. M., Amores, A., Fuiten, A. M., Brown, R. S., … Cresko, W. A. (2016). The genome of the Gulf pipefish enables understanding of evolutionary innovations. Genome Biology, 17(1), 258. doi: 10.1186/s13059-016-1126-6

Smith, H. O., & Welcox, K. W. (1970). A Restriction enzyme from Hemophilus influenzae. Journal of Molecular Biology, 51(2), 379–391. doi: 10.1016/0022-2836(70)90149-X

Stankowski, S., Chase, M. A., Fuiten, A. M., Rodrigues, M. F., Ralph, P. L., & Streisfeld, M. A. (2019). Widespread selection and gene flow shape the genomic landscape during a radiation of monkeyflowers. PLOS Biology, 17(7), e3000391. doi: 10.1371/journal.pbio.3000391

Stuart, Y. E., Veen, T., Weber, J. N., Hanson, D., Ravinet, M., Lohman, B. K., … Bolnick, D. I. (2017). Contrasting effects of environment and genetics generate a continuum of parallel evolution. Nature Ecology & Evolution, 1(6), 0158. doi: 10.1038/s41559-017-0158

Suchan, T., Espíndola, A., Rutschmann, S., Emerson, B. C., Gori, K., Dessimoz, C., … Alvarez, N. (2017). Assessing the potential of RAD-sequencing to resolve phylogenetic relationships within species radiations: The fly genus Chiastocheta (Diptera: Anthomyiidae) as a case study. Molecular Phylogenetics and Evolution, 114, 189–198. doi: 10.1016/j.ympev.2017.06.012

Timm, H., Weigand, H., Weiss, M., Leese, F., & Rahmann, S. (2018). ddrage: A data set generator to evaluate ddRADseq analysis software. Molecular Ecology Resources, 18(3), 681–690. doi: 10.1111/1755-0998.12743

Toonen, R. J., Puritz, J. B., Forsman, Z. H., Whitney, J. L., Fernandez-Silva, I., Andrews, K. R., & Bird, C. E. (2013). ezRAD: a simplified method for genomic genotyping in non-model organisms. PeerJ, 1, e203. doi: 10.7717/peerj.203

Varadharajan, S., Sandve, S. R., Gillard, G. B., Tørresen, O. K., Mulugeta, T. D., Hvidsten, T. R., … Jakobsen, K. S. (2018). The Grayling Genome Reveals Selection on Gene Expression Regulation after Whole-Genome Duplication. Genome Biology and Evolution, 10(10), 2785–2800. doi: 10.1093/gbe/evy201

Wang, S., Meyer, E., McKay, J. K., & Matz, M. V. (2012). 2b-RAD: a simple and flexible method for genome-wide genotyping. Nature Methods, 9(8), 808–810. doi: 10.1038/nmeth.2023

